# Keeping Your Eye, Head, and Hand on the Ball: Rapidly Orchestrated Visuomotor Behavior in a Continuous Action Task

**DOI:** 10.1101/2024.12.17.628893

**Authors:** Anna Schroeger, Alexander Goettker, Doris I. Braun, Karl R. Gegenfurtner

**Author notes:** Correspondence should be addressed to: Anna Schroeger Abteilung Allgemeine Psychologie, Justus Liebig University Giessen, Germany.

## Abstract

In everyday life, we must adapt our behavior to a continuous stream of tasks and time motor responses and periods of resting accordingly. To mimic these challenges, we used a continuous interception computer game (Pong) on an iPad. This allowed us to measure the coordination of eye, hand, and head movements during natural sequential behavior while maintaining the benefits of experimental control. Participants intercepted a moving ball by sliding a paddle at the bottom of the screen so that the ball bounced back and moved toward the computerized opponent. We tested i) how participants adapted their eye, hand, and head movements to this dynamic, continuous task, ii) whether these adaptations are related to interception performance, and iii) how their behavior changed under different conditions and iv) over time. We showed that all movements are carefully adapted to the upcoming action. Pursuit eye movements provide crucial motion information and are emphasized shortly before participants must act; a strategy associated with better performance. Participants also increasingly used pursuit eye movements under more difficult conditions (fast targets and small paddles). Saccades, blinks, and head movements, which would lead to information loss, are minimized at critical times of interception. These strategic patterns are intuitively established and maintained over time and across manipulations. We conclude that humans carefully orchestrate their full repertoire of movements to aid performance and finely adjust them to the changing demands of our environment.

## Introduction

Eye movement measurements have become an important and highly informative behavioral tool to gain insights into active vision during action, planning and decision processes (Gold & Shadlen, 2000; Hayhoe, 2017; Hayhoe & Ballard, 2005; Orquin & Loose, 2013; Schütz et al., 2011; Spering, 2022). Human eyes have a central fovea within their retinae, a small region densely packed with photoreceptors that allows high acuity vision (Osterberg, 1935). Accordingly, the eyes are continuously directed to relevant locations to benefit from the sharp, detailed foveal vision and the corresponding enormous central processing resources (for reviews, see Stewart et al., 2020; Strasburger et al., 2011). Therefore, the analysis of eye movements can provide an objective and implicit ‘window into the mind’ (van Gompel et al., 2007). So far, many studies on the fundamentals of oculomotor control often used a repetitive trial-by-trial structure to measure single reactions to the same artificial target stimulus. While this approach offers the highest level of experimental control to directly relate low-level stimulus properties, like position, speed or contrast to changes in behavior (Becker & Fuchs, 1969; Carl & Gellman, 1987; Goettker, Braun, & Gegenfurtner, 2019; Rashbass, 1961; Spering et al., 2005), it lacks important features of natural behavior (see Miller et al., 2022). Instead of single eye movement responses to the same event we aim to investigate the coordination of eye, head, and hand movements in response to dynamic, sequential task demands that mimic the requirements of our natural environment.

Natural behavior differs in several ways from the typical lab setting (e.g., Foulsham et al., 2011). First, at each moment in time, the real world provides us with rich information that forces us to select and direct our attention to the most relevant object or area (Nuthmann et al., 2010; Tatler et al., 2017). In contrast, in a simplified lab experiment participants often just need to attend to a single and simple object. Second, in the real world we can interact with and change our environment over time (Findlay & Gilchrist, 2003), instead of just passively observing and reacting to sequences of very similar stimuli during large numbers of trials. Third, instead of planning only a single action during a short trial, we usually plan action sequences (Hallett & Lightstone, 1976; Hoppe & Rothkopf, 2019; Najemnik & Geisler, 2005; Renninger et al., 2005; Renninger et al., 2007; Wagner et al., 2023) to achieve a certain goal.

Choosing tasks that impose realistic, less predictable, dynamic demands is essential for investigating naturalistic behavior, because task demands can fundamentally shape how we move (Epelboim, 1998; Goettker et al., 2025; Hayhoe & Ballard, 2005; Rothkopf et al., 2007). For example, in lab paradigms the head is often constrained, but head movements are an integral part of natural behavior in active tasks. There are more and faster head movements when we manually interact with a sequence of targets (tapping) compared to simply observing this sequence (Epelboim, 1998). While eye movements enhance the processing of previously peripheral locations, head movements can serve to explore new areas (Solman et al., 2017). In the context of interacting with moving objects, head movements can also serve to support tracking the object, for example in baseball (Bahill & LaRitz, 1984; Higuchi et al., 2018). The head gets more involved when professional batters track the baseball, compared to when students are at bat (Bahill & LaRitz, 1984). Head movement velocities in baseball players increase shortly before batting (Higuchi et al., 2018). However, head movements can also introduce additional noise due to additionally required sensorimotor transformations (Abedi Khoozani et al., 2020). This might lead to a suppression of these movements shortly before and during actions when tracking can be achieved simply by moving the eyes. Moreover, when a task contains temporal regularities, eye movements are adjusted to sample information (Hoppe & Rothkopf, 2016) and minimize information loss (Hoppe et al., 2018; Shalom et al., 2011) at critical times. Anticipatory eye responses often precede future target movements, indicating that participants strategically time their gaze shifts to align with upcoming events (Diaz et al., 2013; Goettker et al., 2023; Goettker et al., 2021; Hayhoe et al., 2012; for review, see Kowler et al., 2019; Land & McLeod, 2000). Not only the point in time when the eyes move, but also where they move can be adapted to new demands, as demonstrated by saccadic adaptation (Deubel et al., 1986; McLaughlin, 1967; Schütz et al., 2014; Souto et al., 2016). These and similar adaptation processes can provide an ideal vehicle for understanding learning processes (Chen-Harris et al., 2008; Herman et al., 2013; Sailer et al., 2005; Wagner & Schütz, 2023). In summary, understanding how we dynamically adapt our behavior to varying task demands provides valuable insights into human behavior in naturalistic settings.

In this study, we use a gamified, continuous interception task (Pong) on an iPad to examine sequences of simultaneously recorded eye, hand, and head movements. By leveraging the technical developments of mobile eye tracking systems, our setup allowed us to investigate the joint coordination of eye, hand, and head movements without restricting natural behavior through a chin rest. We focus on how these movements are adapted in response to a sequence of actions, across different time scales, and under experimental manipulations of ball speed and paddle size. These manipulations change the overall task demands (i.e., exploring a range of difficulties) to investigate how stable the behavioral patterns are and whether participants can adapt to these new challenges during the game. Our paradigm fulfills several aspects of a naturalistic task while still offering a high level of experimental control. It presents participants with several moving objects they need to attend to: their own paddle on the bottom of the screen, an automated paddle on the top of the screen, and the ball. Instead of passively observing, participants continuously interact with the scene by positioning their own paddle to intercept the ball and thereby shape its future trajectory. The task requires to plan and execute a sequence of actions until one of the paddles misses the ball. We show that behavior is tailored to the intricate dynamic requirements of the task. Across all experimental conditions and over time, pursuit, saccades and blinks, but also head and hand movements are properly timed to keep the ball in foveated vision shortly before intercepting it, while not losing visual information at crucial times. This complex orchestration of behavior is intuitively established and benefits interception performance.

## Methods

### Participants

A total of 24 participants (16 females, 3 left-handed, mean age: 25 (sd = 3.7) years) completed the experiment so that their full data sets are available (2 additional recordings were incomplete and therefore not included in the analysis). For head movements analyses, data of only 22 participants was available due to technical issues during the recording. All participants had self-reported normal or corrected-to-normal vision. Participants gave informed consent prior to participating in the study and received a compensation of 8€ or credits for 1h of participation. This experiment is part of a project approved by the ethics committee of the University of Giessen (approval number: LEK-2021-0028).

### Materials

The Pong Game was presented on an iPad 12 Pro (2732 x 2048 px, 264 ppi). The game consists of a black dot representing the ball (radius: 19 mm = 20 px) and two black rectangular paddles with rounded edges, one controlled by the participant and one automated (the opponent) presented on a uniform white background (screen size: 26.3 cm x 19.7 cm). The opponent’s paddle was programmed to move horizontally with the ball when the ball approached it, but not when the ball moved away. We set the velocity of the physics body of the “opponent” to reach the current horizontal ball position but added a small and variable speed so that it did not perfectly match the ball’s position. Accordingly, the opponent always aimed for the approximate location of the ball. Every 15 actions the opponent was meant to miss the ball, which we achieved by shifting the position the opponent was targeting for with respect to the ball. Instead of aiming for the ball the opponent was aiming for a shifted location during the whole 15^th^ approaching phase. We manipulated paddle width (small: 1.5 cm = 160 px and large: 2.3 cm = 240 px), while keeping the height constant (120 px = 1.15 cm). The ball moved linearly with a constant speed over the screen and bounced back from the side walls and paddles. Between blocks, the ball speed (*slow*: 27 px/frame = 15.6 cm/s ≈ 17.3 deg/s, *fast*: 37 px/frame = 21.4 cm/s ≈ 23.2 deg/s; assuming a distance of approximately 50 cm between eyes and iPad) was changed. All bounce angles except the bounce from the opponent paddle followed physics laws, meaning that the emergent angle equaled the incidence angle. We systematically manipulated the bounce angle from the opponent paddle to achieve variable ball trajectories and control the angle in which the ball approached the participant’s paddle. We kept the natural bounce direction but adapted the size of the angle between the perpendicular and the ball direction after the bounce randomly between 15, 45, or 75 degrees. The largest angle led to several bounces between the walls until the ball reached the participant, whilst the smallest angle led to direct trajectories without any or with only a few ball bounces.

Eye and head movements were recorded with Tobii Glasses Pro 3 and analyzed in Matlab (The MathWorks Inc., 2022) using the GlassesViewer (Niehorster et al., 2020) and self-written code. The sampling rate of gaze data was 100 Hz, scene camera 25 Hz, and the IMU ∼100 Hz. The iPad displayed stimuli and saved paddle and ball positions with 60 Hz. For the analysis, all data was interpolated to 100 Hz.

### Procedure

Prior to playing the game, participants were asked for their age, handedness, and whether they were wearing contact lenses. The examiner instructed them to use their index finger by showing them how to slide the paddle horizontally. They were told to intercept the ball but were not given any instructions on how to achieve that. Participants used their dominant hand to play the game, except for one left-handed participant who switched hands occasionally. Three participants switched towards using the middle finger during the experiment. The experiment consisted of four experimental blocks (2 ball speeds x 2 paddle sizes, see **Fig. 1F**) of at least 60 interception attempts, each preceded by a baseline of at least 15 interception attempts with the larger paddle size and slower ball speed. As soon as the minimal number of interceptions for each condition or baseline was reached, the switch to the next condition occurred during the short break following one player scoring a point. The order of the four blocks was counterbalanced across participants. After either the participant or the opponent missed the ball, a fixation cross was presented at the center of the screen for a random interval between 2.5-3 s (uniform distribution). Subsequently, the ball started moving linearly from the center toward the opponent. Before starting the experiment participants were familiarized with the mobile eye tracker and a calibration and validation were done (glassesValidator, Niehorster et al., 2023). The validation was repeated after the experiment to ensure sufficiently accurate data quality throughout the time of experiment. Altogether the whole experimental session took about 1h.

**Figure 1.**
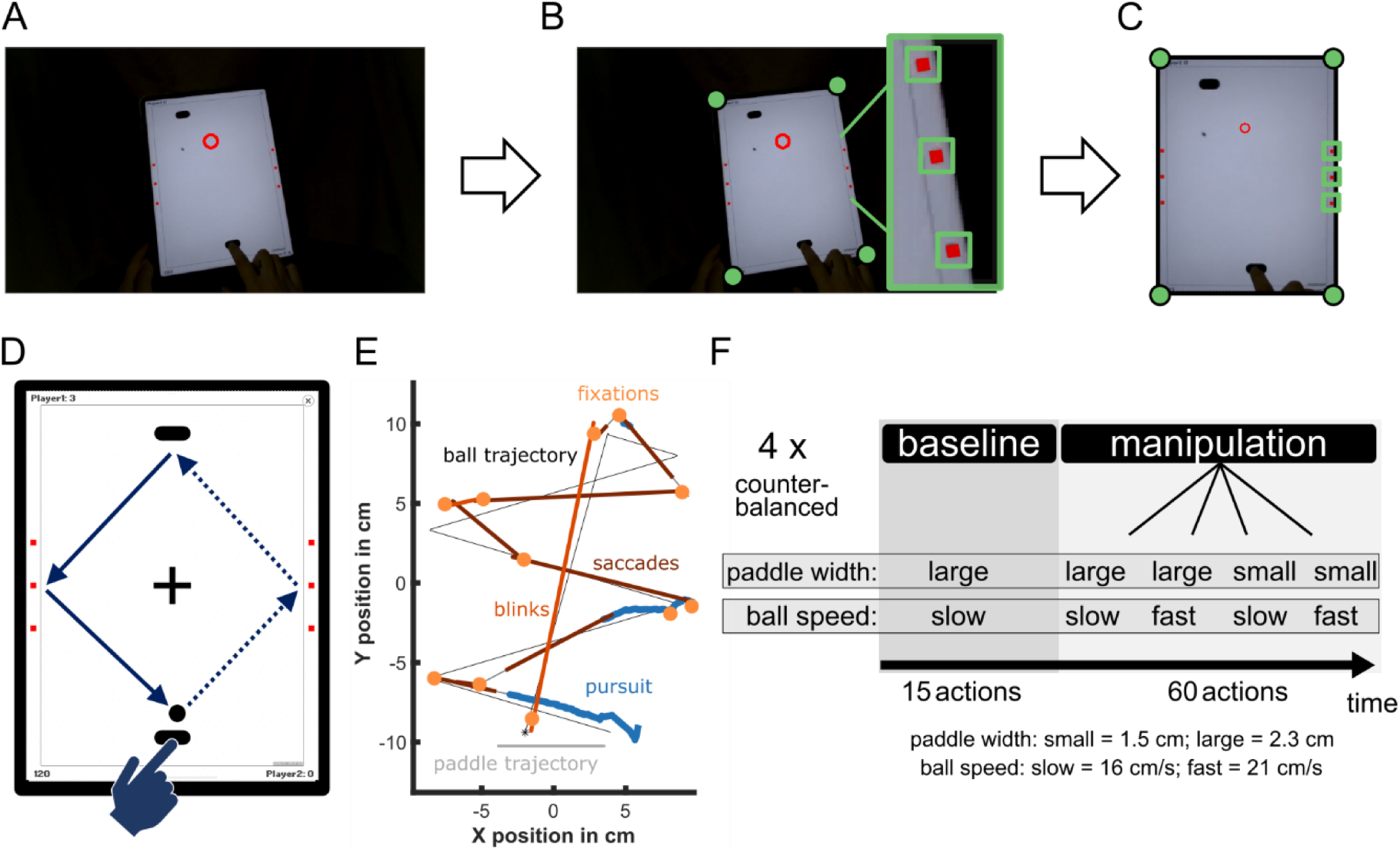
Illustration of the video game Pong and the methods. A) View of the iPad through the front camera of the mobile eye-tracker with the gaze position indicated by a red circle. B) We detected the red reference markers (three red squares on each side) or the corners of the iPad (illustrated by green dots) in a binary version of each frame using a color mask and/or luminance threshold. C) A homography transformation aligns the image and the gaze coordinates to the iPad’s dimensions. D) Illustration of an action cycle: starting with the participant’s interception (bottom), followed by the return of the opponent (top) and concluding with the next interception. The task demands largely differ between the phase in which the ball recedes from the participant’s paddle (dotted line) compared to the phase in which the ball is approaching (solid line). Therefore, we expect differences in gaze behavior across the time of an action cycle. E) The ball trajectory (black line, start marked with an asterisk) and gaze positions are shown for a representative action cycle. Participants track the ball using a mixture of saccades (red segments) and pursuit (blue segments) with fixations (yellow circles) primarily near bounce points. Gaze positions also change during blinks (orange segments). In this example, the ball moves directly towards the opponent without wall bounces, is returned, and bounces twice on each side before reaching the paddle of the participant. F) Experimental Procedure: participants completed four manipulation blocks, each preceded by a baseline of 15 interception attempts with the slower ball and larger paddle.

### Eye and Head Movement Analysis

In a first step the gaze data was manually temporally aligned with the iPad using the scene video and a green rectangle displayed on the screen at the beginning of the game only for this purpose. The gaze position within each video frame was then transformed to the iPad’s reference frame by using computer vision algorithms in MATLAB version R2022a (The MathWorks Inc., 2022). For each video frame of the scene camera, the position of the iPad was detected using a color mask and object detection with the MATLAB function bwboundaries. If possible, this was done by detecting the position of six red rectangular reference objects (see **Fig. 1B**). If this procedure did not result in detecting six objects, we simply ran bwboundaries on a binary version of the frame to detect the outline of the screen and extracted the position of the four corners. With either the six or four detected positions we performed a homography transformation (see **Fig. 1**) using Matlab functions fitgeotrans (projective) and imwarp. This procedure allows us to remap gaze positions within each video frame to the actual position on the iPad’s screen. We filtered the gaze positions and eye velocity signal (both 100 Hz) using a third-order Savitzky-Golay filter of frame length 11 (110ms).

Using the original eye movement traces before transformation, saccades and blinks (missing data episodes) were classified as implemented in the glassesViewer (Niehorster et al., 2020) choosing the adaptive velocity algorithm from Hessels et al. (2020). In a second step, we relabeled saccades with durations equal to or longer than 200 ms as blinks (Gandhi & Katnani, 2012; Leigh & Zee, 2015). Third, based on the gaze position relative to the iPad, pursuit events were detected: every episode lasting at least 100 ms with a gaze velocity of at least 7 cm/s (∼50% of the slower ball velocity), and moving in the same direction as of one of the three objects, the ball and the two paddles (+-30 degrees). Each pursuit event was related to exactly one of these targets, namely the target with the most similar movement direction. All episodes of at least 100 ms in which the eyes did not fulfill any of the other criteria (= nearly no motion in gaze position) were labeled as fixations. Episodes shorter than 100 ms which were not attributed to one of the labels and episodes with missing gaze signal were not labeled as eye movements (16 %).

To detect head movements, the gyroscope data of the eye tracker was used, and all three velocity directions were combined to one measure: the velocity of the total movement (square-root of the summed squared velocities), which was filtered in the same way as the eye velocities. Times in which the overall velocity was at least 10 deg/s for at least 100 ms were labeled as head movement events. Head direction angles were computed from the directional gyroscope data of the head mounted eye-tracker and evaluated relative to the target motion direction.

The horizontal position of the paddle, recorded by the iPad, served to extract hand movements. The signal was first interpolated to 100 Hz and then filtered with a third-order Savitzky-Golay filter with a frame length of 11 (110 ms). We detected hand movement intervals in the signal, when the velocity exceeded 5cm/s for at least 100 ms.

### Statistical Analysis

To analyze the adaptation of eye movements over time only complete episodes of an action cycle from one interception (or the start from the center) to the next interception were included. The frame after the ball bounced from the participants’ paddle was labeled as the start frame and the frame of the next bounce from the participant’s paddle was labeled as end frame. In between the ball bounced once from the opponent. If more than 45% of the gaze data between those frames were missing, the whole episode was excluded.

We summarized the proportions of eye, head and hand movements per participant and point in time (relative to the next action) in steps of 0.1%. Difference-scores were calculated for each participant. For statistical analyses we used R (R Core Team, 2021) and RStudio (RStudio Team, 2016). To test for aggregated differences between phases, between hits or misses or over time, we report means and t-tests. If the data violated the assumption of normal distribution additional non-parametric Wilcoxon signed rank exact tests were executed, but they revealed the same results. To test for overall effects of the two manipulations, repeated measures ANOVAs (package ez, Lawrence, 2016) on the performance or summarized gaze statistics were performed using the within-subject factors paddle size (small or large) and ball speed (slow or fast). To contrast successful interceptions with misses and analyze the exact time windows, we fitted Generalized Additive Models (package itsadug, van Rij et al., 2020; package mgcv, Wood, 2011) including either random intercepts, both random intercepts and random slopes or random intercepts and random smooths to the data streams. For this purpose, we resampled the data to reflect the actual sampling rate (instead of steps of 0.1% relative time, the steps were chosen based on the average duration of an action cycle). Using *plot_diff* we derived significant intervals for differences between hit and miss actions.

## Results

While playing Pong, participants moved their eyes to mostly track the ball, the opponent’s paddle, or their own paddle (see **Fig. 1E**). Only occasionally gaze was directed toward the players’ scores or off-screen (see **Table 1** and **Fig. 2**). **Table 1** shows that we observed all sorts of eye movements during the game: pursuit, saccades, fixations, and blinks. **Fig. 2** highlights the importance of the relative timing of eye, hand and head movements.

**Figure 2.**
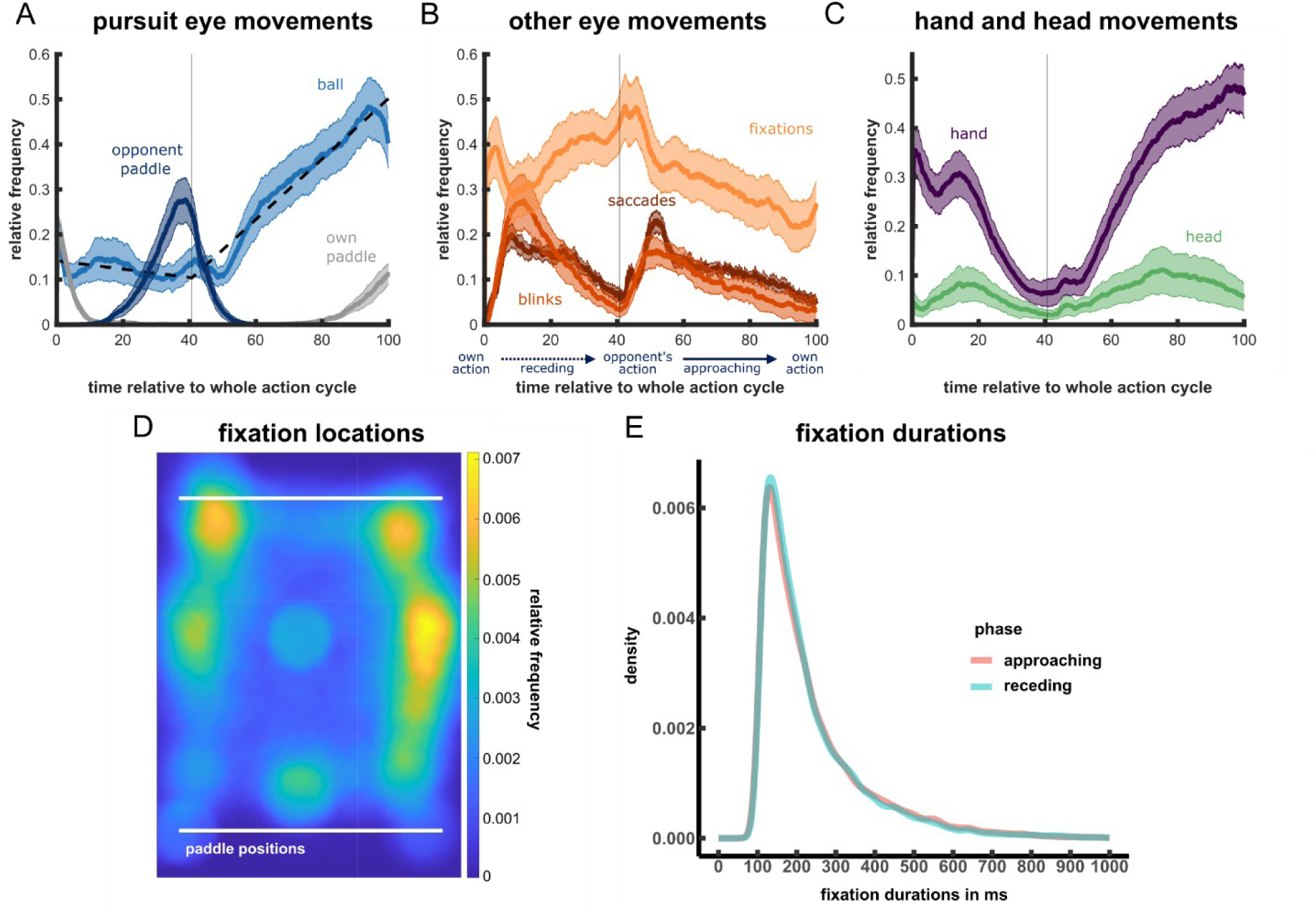
Frequencies of the different eye movement types, hand movements, head movements, and bounces (mean and 95% between-subject intervals) over time during the action cycles. Relative frequencies depict the proportion of action cycles in which the respective behavior was shown at each time point. The vertical line at 41% in A-C illustrates the moment the opponent acted. A) The choice of the pursuit target changed during the action cycle depending on the ball position: participants pursued either paddle movement (own: grey, opponent: dark blue) when the ball was close to one of them. Pursuit of the ball (bright blue) steadily increased during the approaching but not during the receding phase. The dashed line indicates a 2-piece linear fit with the two pieces crossing around the action of the opponent. B) Fixations (bright orange) become less frequent with the approach of the next action, blinks (medium orange), and saccades (dark orange) are reduced during the opponent’s and participant’s actions. C) Hand movements (dark purple) to control the paddle were present during interception and were largely reduced around the opponent’s action. Head movements (bright green) were less frequent than eye movements. Participants moved their heads mainly before intercepting the ball and these movements were more closely aligned with the ball trajectory compared to early head movements. Please note, that in ∼16% per action cycle gaze behavior did not fulfill any criterion and was therefore not classified as any of these events (see Methods for more details). Consequently, there are non-classified episodes and overall, the curves in Fig. 2A and B do not add up to 1. D) Fixation locations across all participants. E) Fixation durations in receding vs. approaching phase.

**Table 1.** Descriptive statistics of the proportion of samples labeled as each gaze status averaged over an action cycle. Please note that only gaze positions on the iPad are considered and gaze off-screen as well as missing data are excluded.

*Descriptive statistics on Proportions of eye movements as percentages*
| Variable | Mean | SD | Min | Max |
| --- | --- | --- | --- | --- |
| Pursuit ball | 23.83 | 7.25 | 13.99 | 42.70 |
| Pursuit own paddle | 2.00 | 0.83 | 0.03 | 3.83 |
| Pursuit opponent | 4.36 | 2.12 | 0.00 | 8.67 |
| Saccades | 14.02 | 1.60 | 10.89 | 17.72 |
| Fixations | 34.76 | 12.01 | 3.22 | 55.91 |
| Blinks | 6.48 | 5.03 | 0.29 | 14.82 |

Our setup allowed us to investigate how gaze behavior was adapted across multiple time scales and whether these adaptations benefit game performance: First, we tested whether and how eye, head and hand movements changed across the continuous sequence of the ball bouncing from the participant’s paddle to the opponent’s paddle and back. Second, we examined how differences in behavior were related to task performance. Third, we analyzed how behavior generally adapted through experience over playing time, and fourth, across varying demands of ball speed and paddle size.

### Do we adapt to the dynamic demands of the task?

**Fig. 2 and Table 1** illustrate the relative frequency of the investigated behavior across the time of an action cycle: pursuit (**Fig. 2A**), saccades, blinks and fixations (**Fig. 2B**), and hand and head movements (**Fig. 2C**). Do we adapt the type and timing of movements to the dynamic demands of the task? We analyzed how frequent each behavior was at each moment in time relative to the own action. We normalized the time between one interception of the participant and the next interception with one bounce from the opponent in between. The duration of one typical action cycle (see **Fig. 1D**) ranged between 1.2 s and 9.3 s (95% of durations, *median* = 3.4 s). The receding phase, in which the ball moved from the participant towards the opponent, was on average 1.7s long, while the approaching phase took 2.5s. This difference in durations is also illustrated in **Fig. 2A-C**, where the bounce of the opponent is located around 41% of relative time.

The most striking effect was on the frequency of pursuit eye movements: the closer the ball approached the participant’s paddle the more likely the participant pursued the ball – that is, they smoothly tracked the ball to keep it foveated (bright blue line in **Fig. 2A**). Fitting a 2-piece linear function to the time course indicated that the increase in pursuit frequency started shortly after the ball was hit by the opponent’s paddle (see dashed lines in **Fig. 2A**). When comparing the slopes, the increase in ball pursuit during the approaching phase is significant, *t*(23) = 5.88, *p* < .001, *d* = 1.200. In contrast, participants tracked the paddle of the opponent instead of the ball when the ball was close to it (∼27% - 44% of the time, dark blue line in **Fig 2A**). While pursuit of the ball increased in the second half, the frequency of fixations diminished (bright orange line in **Fig. 2B**). Participants rarely tracked their own paddle, which was controlled by their own hand movement.

During and shortly before the ball hit one of the paddles (41% and 100% of relative time) participants rarely blinked (medium orange line in **Fig. 2B**). Instead, participants seemed to blink mainly during time periods shortly after the ball bounced back from their own paddle or from the opponent’s paddle (vertical line in **Fig. 2B**), so at time periods with the least critical visual information. In line with this, participants also blinked more frequently in time periods after they had hit the ball with their paddle, and a lot of time was still left until they had to intercept the ball again, compared to when the opponent returned the ball and it was approaching them (mean of 0.15 vs. 0.11 blinks per second), *t*(23) = 2.99, *p* = .007, *d* = 0.610. Additionally, blinks were initiated earlier in receding compared to approaching phases (mean of 0.86 s vs. 1.26 s), *t*(23) = 2.67, *p* = .014, *d* = 0.544. Please note that *relative* blink initiation time was equal between phases (31 % vs. 32 % of relative time, *p* = .810), which is caused by different mean durations of the phases (receding: 1.7 s, approaching: 2.5 s).

Saccades showed a similar time course as blinks (dark orange line in **Fig. 2B**): Most saccades occurred during the middle of the receding and approaching phases. Relative to the start of each phase, saccades were executed on average ∼143 ms later in the approaching phase compared to the receding phase (mean of 1.21 s vs. 1.06 s), *t*(23) = 3.37, *p* = .003, *d* = 0.687. Please note that the *relative* timing was earlier in the approaching phase due to different phase durations (49% vs. 45 % of relative time, *p* < .001). Saccade rates (count per second) were not significantly different between phases (*p* = .364).

Participants used fixations throughout the action cycle, but fixation frequency diminished as the action approached (**Fig. 2B**). Fixations occurred mostly along the ball’s trajectory (see example in **Fig. 1E**) and mainly focused on the walls and the paddles (**Fig. 2D**). Fixation durations showed similar distributions between receding and approaching phase **Fig. 2E**).

Hand movements mainly occurred towards the end of an action cycle or shortly after an action (purple line in **Fig 2C**). Participants seemed to wait for the target while the opponent was interacting with it. Head movements were generally less frequent (green line in **Fig. 2C**) but showed small peaks in the middle of receding and approaching phases. We additionally analyzed whether the direction in which the head pointed changed with the position of the ball. To do so, we compared the angle of the ball’s trajectory during the head movements with the angle of head rotation. If the head pointed towards the ball and perfectly rotated with the ball, the difference between those angles was 0 degrees. These analyses showed that head movements were more closely aligned with the ball trajectory, when the ball was approaching the participant compared to the receding phase (mean difference in direction of 26 vs. 53 deg), *t*(21) = 6.36, *p* < .001, *d* = 1.36.

Together, these results demonstrate that participants specifically adapt their behavior to the changing requirements of the different phases of the dynamic task: Participants try to blink, move their head, and make most of their saccades during times when the ball just started receding from their own or the opponent’s paddle. The closer the moment of interception, the more participants focus on pursuing the ball and aligning their head movements with it.

### Association between eye movements and performance

The above results raise the question whether this typical pattern of movements benefits performance. To test this, we analyzed behavior, especially pursuit eye movements, with respect to hit vs. miss actions within observers, but also across observers. Within participants, Generalized Additive Mixed Models (GAMMs) fitted to ball pursuit frequencies over time revealed significant differences between hits and misses. Significant intervals are indicated by bars at the bottom of **Figure 3A**. If a participant hit the ball in a given action cycle, it was more likely that the preceding eye movement was pursuit compared to action cycles in which they missed it (**Fig. 3A**). The fact that the general increase in ball pursuit during the approaching phase (see **Fig. 2A**) was even larger before a hit highlights the specific importance of ball pursuit during the final phase before interception. In contrast, early pursuit in each action cycle, especially shortly after the bounce at the opponent’s paddle was associated with missing the ball. Additional analyses revealed that more hand and head movements towards the end of the action cycle, as well as pursuit of the own paddle were related to worse performance, as indicated by the significant intervals of the GAMMs (**Fig. 3A and B**).

**Figure 3.**
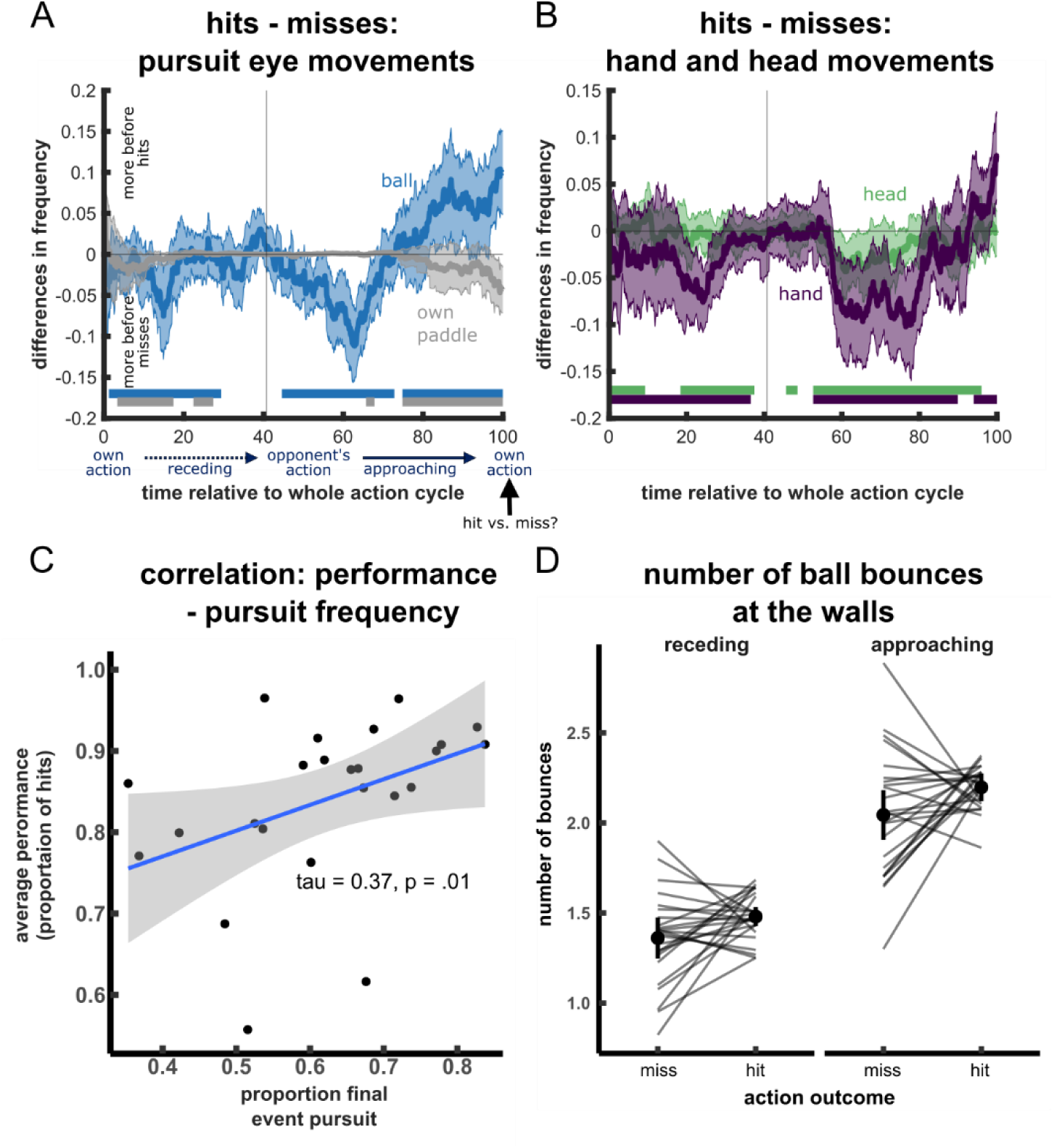
Association between performance and eye, hand, and head movements. A-B) Differences in Eye, Head and Hand Movement Frequencies for hit versus miss actions. Shown are the means and 95% between-subject confidence intervals. Positive values indicate that the respective movement was more frequent prior to hit actions at a specific point in time. Generalized additive mixed models were fitted to the data and time intervals of significant differences between hits and misses are indicated by the bars at the bottom. Pursuit of the ball (bright blue) shortly before interception was associated with hitting the ball, while shortly after the opponent’s action it was rather associated with missing it. Shortly before missing the ball participants pursued the movements of their own paddle (bright blue) more frequently compared to the same time when hitting the ball. Hand (dark purple) and head (bright green) movements shortly before interception were indicative of missing the ball. C) Between-subject correlation between pursuit and performance. Participants who show more frequently pursuit as a final eye movement before hitting the ball, perform better in the game. D) Number of ball bounces in receding vs. approaching phases for successful hits and misses.

The beneficial effect of ball pursuit was further supported by analyses across participants. Participants who were more likely to pursue the ball movement immediately before or during interception scored higher in the game (**Fig. 3C**), τ = 0.37, *p* = .01. In summary, this indicates that the movement patterns shortly before acting are indicative of the interception success. Specifically, pursuit of the ball movement before interception improves interception performance.

Given that pursuit typically stops when the ball bounces, the number of bounces might cause differences in pursuit frequencies. The ball bounced off the side walls on average slightly more frequently preceding hits (3.7 bounces) compared to misses (3.4 bounces), *F*(1, 23) = 8.93, *p* < .007, η_p_^2^ = .28. It also bounced more frequently in the approaching (2.1 bounces) compared to the receding phase (1.4 bounces), *F*(1, 23) = 278.55, *p* < .001, η_p_^2^ = .92; **Fig 3D**). We therefore compared the distribution of bounce timings in the approaching phase between hit and miss trials. According to separate Kolmogorov-Smirnov-Tests for each participant, the bounce timing did not differ between hit and miss trials for 15 out of 24 participants (mean *p*-value = .190). This suggests that the differences in pursuit frequency between successful actions and misses cannot be simply explained by differences in the timing of bounces.

### Practice related changes

Since practice is known to modify movements patterns, we compared the first 15 action cycles of the initial baseline (preceding the first block) with the 15 action cycles of the final baseline (preceding the final block). Indeed, interception performance increased from 85% to 94% hit rate in later actions (**Fig. 4C**), *t*(21) = 3.26, *p* = .003, *d* = 0.666. Given the importance of pursuit eye movements for interception (e.g., de la Malla et al., 2017; Fooken et al., 2016), we focused on the frequency of ball pursuit (detailed visualizations for other eye movement metrics are provided in **Fig. 5**). Pursuit of the ball did not significantly change over the time of the experiment in the approaching phase (**Fig. 4A**), *t*(23) = -0.50, *p* = .620, d = -0.103. There was a trend for more head movements (**Fig. 4B**), *t*(21) = 2.27, *p* = .033, d = 0.463, but this was not significant after correction for multiple comparisons. Please note that visual inspection of the time course data (see **Fig. 5**) indicates that these effects might be more restricted to specific time windows and therefore are underestimated in the current analysis. It additionally suggests that participants increase pursuit of the opponent over the course of the experiment and slightly increase blink rates at times shortly after opponent actions (see **Fig. 5**). Overall, these analyses show that the movement patterns were intuitively established and relatively stable over the time of the experiment. If behavior was adapted, these adaptations were small and seemed to be limited to specific time windows indicating a fine-tuning of the movement timing instead of a general in- or decrease in movement frequency.

**Figure 4.**
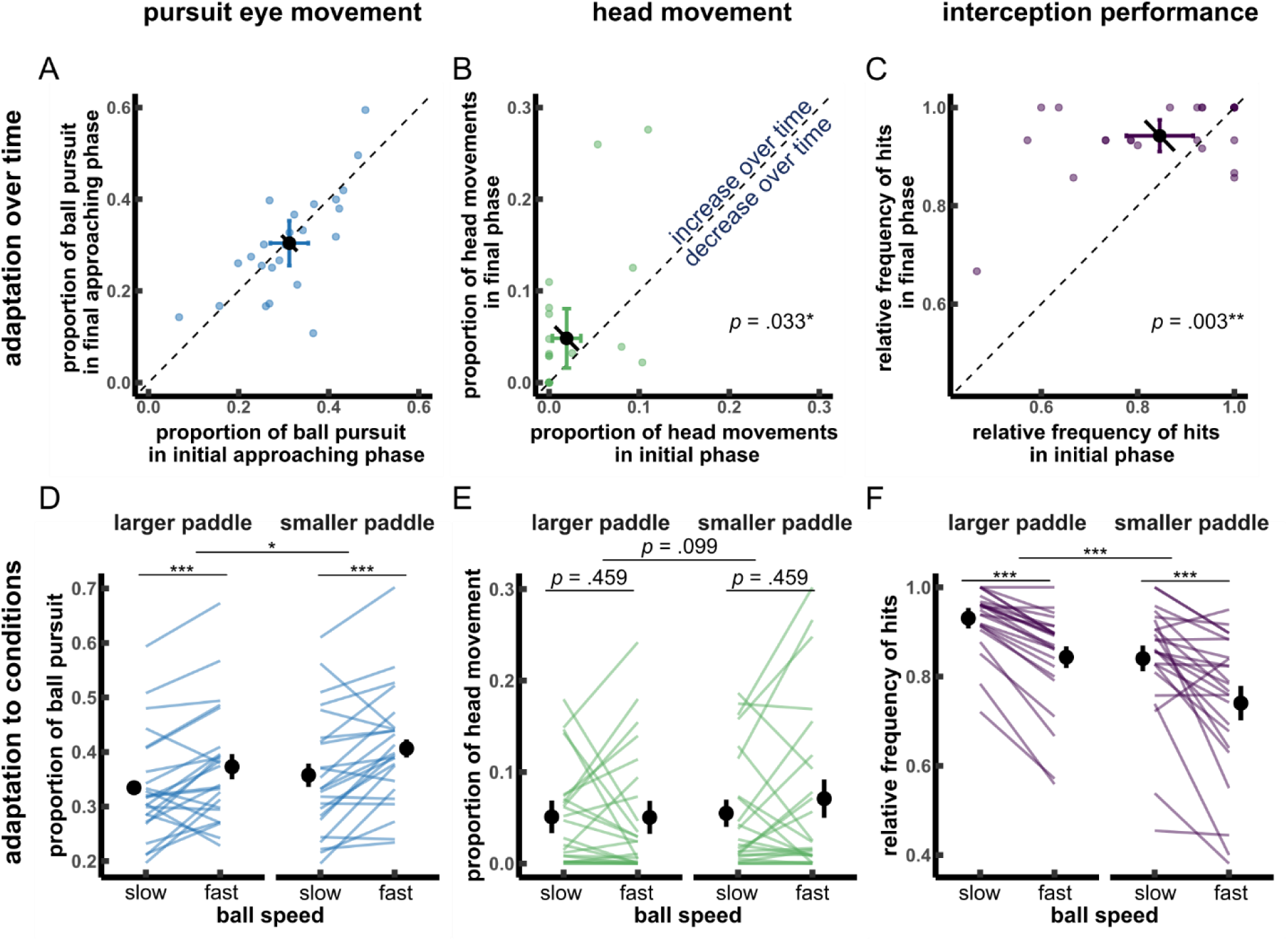
Overview of adaptation effects. Top row: adaptation effects over time of approaching ball pursuit (A), head movements (B) and interception performance (C). Bottom row: adaptation effects of paddle size and ball speed on ball pursuit (D), head movements (E) and interception performance (F). All plots include individual data (colored dots and connecting lines) and means with 95% within-subject confidence intervals (black). The pairwise comparisons for adaptation over time additionally show the uncorrected confidence intervals (colored).

**Figure 5.**
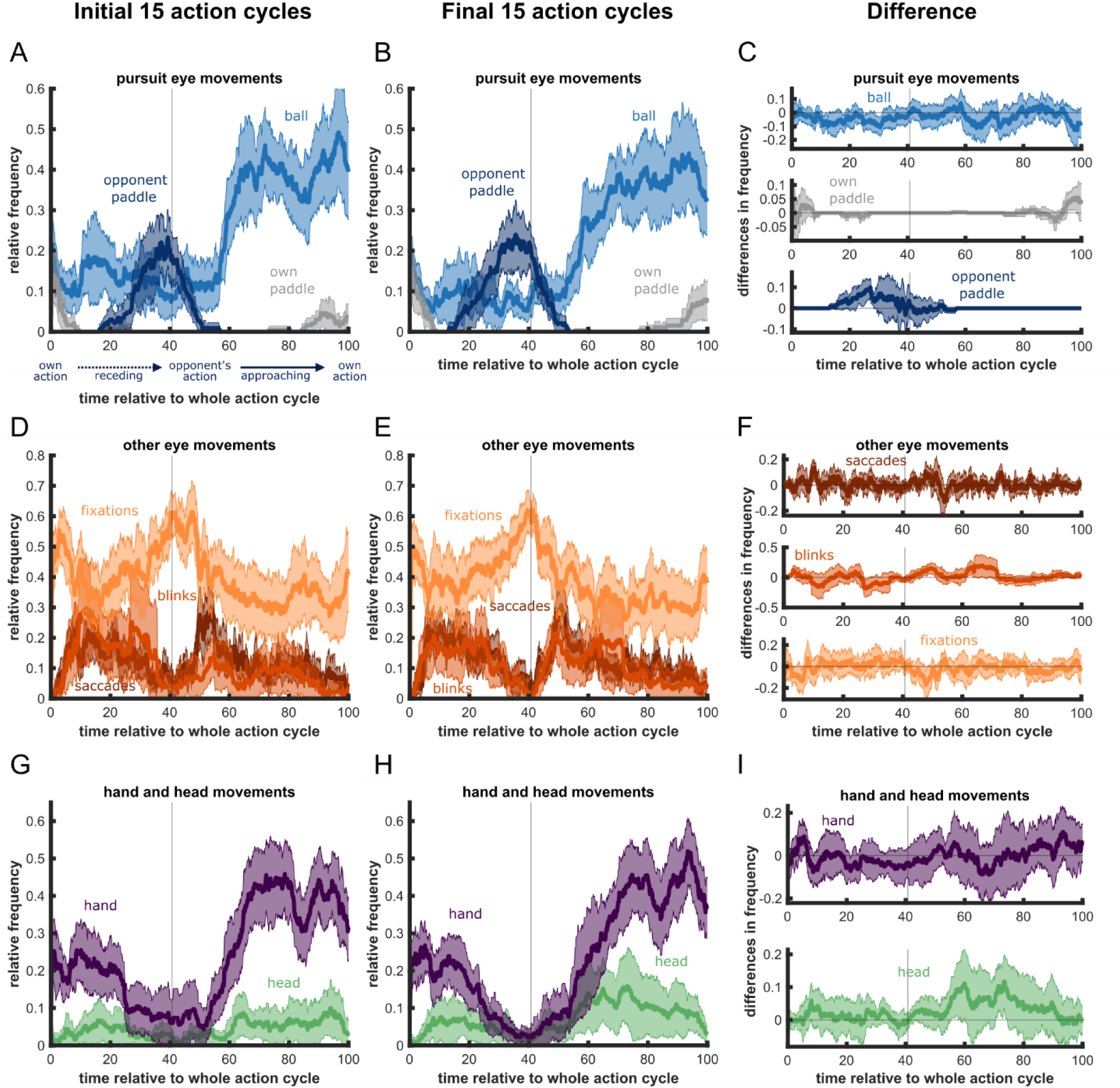
Detailed visualization of adaptation over time. The first column shows eye, head and hand movements for the initial baseline (15 actions) and the second column for the baseline preceding the final block (15 actions). The third column shows differences between early and late actions, with positive values indicating higher frequencies in late action cycles. Overall, the behavioral patterns in the initial (A, D, and G) and final action cycles (B, E, and H) are very similar and resemble the average patterns shown in Fig. 2. Participants do not largely change their strategies. We specifically analyzed potential changes in ball pursuit (C upper panel, approaching phase) and head movements (I lower panel) which are additionally shown in Fig. 4.

### Effect of task demands across experimental conditions

We tested whether ball speed or paddle size impact performance and oculomotor strategies in repeated measures ANOVAs. The overall pattern of eye movements stayed consistent with only small differences between conditions (see **Fig. 6** for detailed visualizations), but there was some fine-tuning. There was more pursuit for faster targets (**Fig. 4D**), *F*(1, 23) = 18.34, *p* < .001, η_p_^2^ = .44, and for the smaller paddle size (**Fig. 4D**), *F*(1, 23) = 9.23, *p* = .006, η_p_^2^ = .29. Head movements were unaffected by ball speed or paddle size (all *p*s ≥ .099, **Fig. 4E**). As expected, interception performance decreased with higher ball speed and smaller paddles (see **Fig. 4F**), as shown by significant main effects of ball speed, *F*(1, 23) = 42.79, *p* < .001, η_p_^2^ = .65 and paddle size, *F*(1, 23) = 35.27, *p* < .001, η_p_^2^ = .61. This indicates that increased pursuit frequency may compensate partially for performance challenges imposed by increased task demands, though head movements showed no significant adaptation.

**Figure 6.**
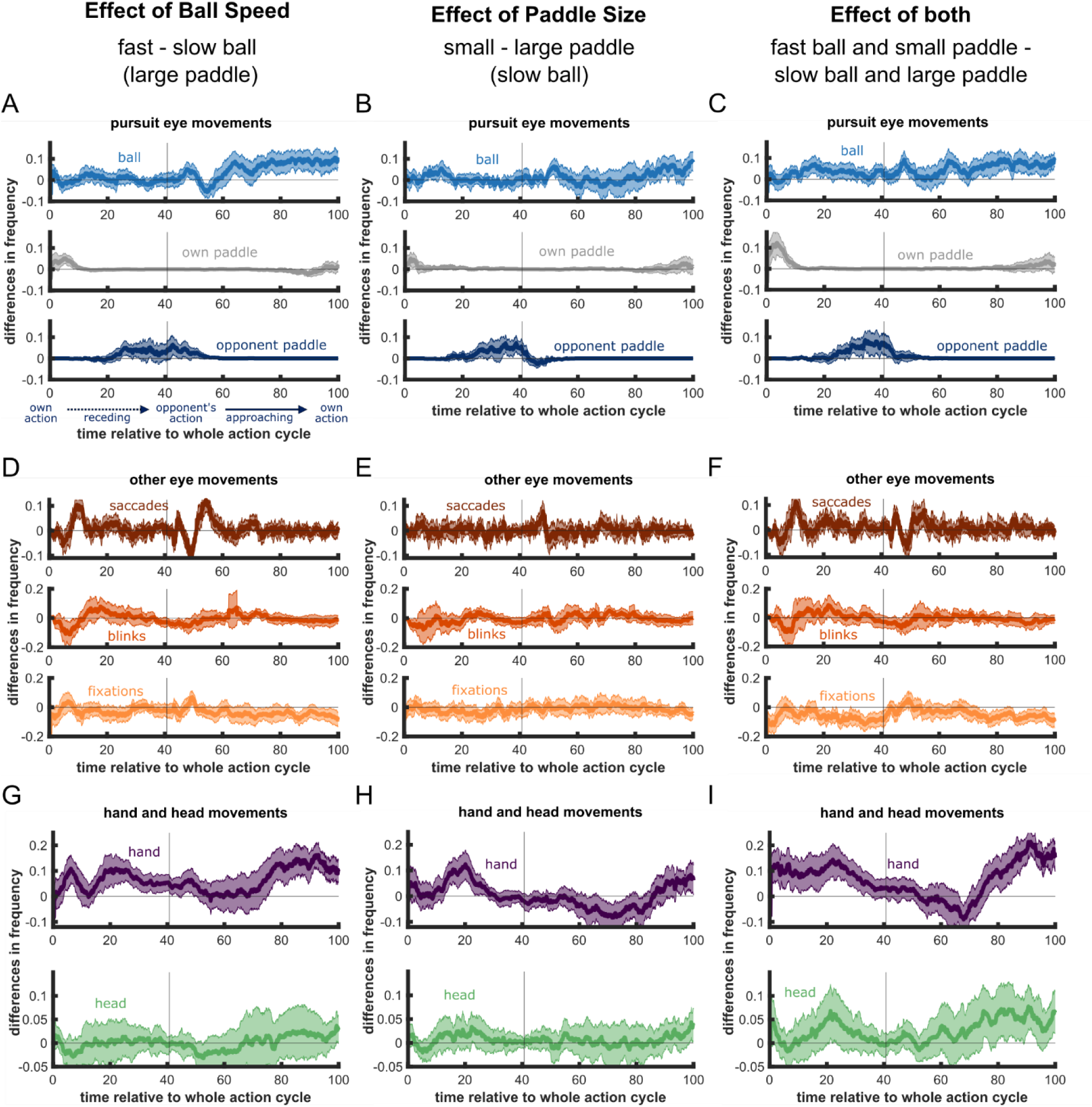
Detailed visualization of adaptation across manipulations. All conditions are compared to the same reference condition with the small paddle and slow ball (same as baseline) and only difference plots to this baseline are shown. The first column indicates the difference between fast and slow ball speeds, the second column indicates the difference between small and large paddles, and the third column shows the differences between the condition of fast ball speed and small paddles with the baseline of slow ball speed and large paddles. Ball pursuit appears to increase for fast balls and small paddles (A, B, and C, upper panel). We analyzed these effects in Fig. 4D and focused on potential changes in head movements in Fig. 4E. Changes in hand movement timing (D, H, I, lower panel) are reflected in performance changes in Fig. 4F. Visual inspection additionally suggests increased pursuit of the opponent for fast balls and small paddles (A, B, C, lower panel). Differences in saccades and blinks across conditions appear small and unsystematic. The increase in ball pursuit seems to come at the cost of slightly decreased fixations (D, E, F, lower panel).

## Discussion

We investigated how participants used and adapted their eye, head, and hand movements for successful performance in a continuous interception task. Our findings indicate that participants developed a robust and effective strategy for intercepting the ball. They carefully orchestrated their eye, head, and hand movements in anticipation of upcoming actions, and this well-coordinated behavior led to more successful outcomes. Participants relied on a blend of saccades and pursuit to track the ball, and specifically increased pursuit in the moments before interception. By contrast, they seldom pursued their own or the opponent’s paddle and minimized potentially distracting movements—such as head shifts, blinks, and saccades—around the critical action moments. Although these patterns remained remarkably stable, participants did make fine adjustments to their movement patterns with practice and under more demanding conditions, suggesting that they refined their strategies rather than overhauling them.

### Adaptation to the action

Participants dynamically adjusted their instrumentation of different motor behaviors to achieve optimal action outcomes. In particular, changes in the proportion of pursuit were a strong indication of the strategic control of eye movements. Firstly, participants increased pursuit frequency as the ball approached their paddle. Secondly, there were significant within-subject differences in the timing of pursuit in relation to successes and failures. Thirdly, there was a significant between-subject correlation between final pursuit frequency and overall game performance. While previous experimental manipulations showed a general advantage of pursuit eye movements compared to fixations for motion perception and interception (de la Malla et al., 2017; for review see Fooken et al., 2021; Spering et al., 2011), we revealed the temporal integration of pursuit within a continuous sequence of actions: Participants intuitively applied but temporally restricted their pursuit to relevant periods in time, maintaining its beneficial outcomes. Shalom et al. (2011) showed that participants kept their eyes closer to the ball before compared to after their interception in a comparable game (‘Breakout’). Our results suggest that this is achieved by closely pursuing the ball before interception. First indications of the great importance of pursuit eye movements were shown in real-world interception, for example by cricket batsman (Land & McLeod, 2000) and squash players (Hayhoe et al., 2012). Our findings on pursuit eye movements contrast with a study using a 2D-interception task with unpredictably moving targets and self-paced timing of interception (Mrotek & Soechting, 2007). In this study, participants smoothly pursued the unpredictable target throughout the whole trial, instead of limiting pursuit to the final part of the trajectory. The reason for this discrepancy might lie in the different temporal demands. When the moment of interception is set as in Pong, participants can focus pursuit to the critical moment, but if they must choose the time of interception themselves, it is relevant to keep pursuing throughout the trial until the right moment is found. However, as in their study, we also observe that saccades are largely reduced during the moment of interception. The positive effect of pursuit for interception might be due to additional information provided by an efference copy (for a review, see Bridgeman, 1995; Sommer & Wurtz, 2008). Thus, the way we track a moving target is predictive of whether we successfully intercept it later. Previous studies have reported that pursuit eye movement behavior in 2D (Brenner & Smeets, 2011; Cesqui et al., 2015; Delle Monache et al., 2015; Fooken & Spering, 2020; Fooken et al., 2016; Kreyenmeier et al., 2017; Spering et al., 2011), and in 3D interception (Arthur et al., 2023; Binaee & Diaz, 2019) can be directly related to performance in such tasks (for review, see Fooken et al., 2021). Here we expand on these findings by showing that participants strategically adjust when to pursue a relevant object even with continuously changing task demands.

We also found that the strategic deployment of different types of movements was not limited to pursuit. The impeding action also affected the timing of blinks, saccades and head movements. Participants restricted behavior that reduces the quality of the visual input to periods that were not critical for game performance (Goettker, Brenner, et al., 2019; Hoppe et al., 2018; Mrotek & Soechting, 2007; Shalom et al., 2011). This is interesting due to three reasons: First, saccades during tracking help to correct position mismatches between the eyes and the target (see Goettker & Gegenfurtner, 2021), which should be beneficial. However, visual sensitivity is reduced during saccades (see Binda & Morrone, 2018; Braun et al., 2021). To manage this tradeoff, participants restricted saccades to moments before interceptions during the action cycle (see **Fig. 2B**). This strategy aligns with findings from an earlier study showing that corrective saccades during tracking are often triggered earlier to avoid that they occur too close to the interception time (Goettker, Brenner, et al., 2019). Similarly, Mrotek and Soechting (2007) found that saccades are largely reduced as soon as an interceptive hand movement was initiated. Our visualizations of an exemplary action cycle and the distribution of fixation locations suggest that participants fixated at predicted target locations mainly along the walls (see **Fig. 1E** and **2E**). They would then continue tracking after the ball bounced, potentially using pursuit eye movements. Similar behavior has been described for 3D interception tasks (Diaz et al., 2013; Hayhoe et al., 2012). Second, even though blinks might have a beneficial effect in specific detection tasks (Yang et al., 2024), our results show that blinks are suppressed at critical times in a dynamic task with motor responses to moving objects. Third, the execution of head movements typically supports visual information intake (Bischof et al., 2023). Previous studies on monkeys (Arora et al., 2019) and humans (Bischof et al., 2024; Burlingham et al., 2024; Hooge et al., 2024) showed that head movements often follow eye movements and serve to center the gaze in the visual field. However, head movements also introduce noise due to the transformation of the visual reference frame (Abedi Khoozani et al., 2020). This was reflected in the fact that head movements toward the end of the trial were related to worse performance. Therefore, during the Pong game participants might have specifically centered their head on the interception location before the critical moment of interception but avoided further head movements during the action. This finding contrasts with previous results from baseball batting which have suggested a supportive role of head movements during tracking (Bahill & LaRitz, 1984; Higuchi et al., 2018). Compared to the ball speeds used in the current paradigm, baseballs move at very high speeds which exceed the maximal possible smooth pursuit speed. This difference might explain why head movements support tracking and potentially also interception performance for fast-paced balls but still hamper performance in the current game with much slower ball speeds where eye movements alone are sufficient to keep track of the target. Overall, the result on head movement patterns again underlines the strategic timing of movements to optimally perform in this task.

In summary, all movement frequencies showed a strong dependency of their timing relative to the execution of interceptive actions, whether by the participant or the opponent. Movements that potentially cause noise or loss of critical visual information, like blinks, saccades and head movements are largely reduced at these critical points in time. In contrast, pursuit eye movements are increased, as they provide additional motion information of direction and speed by efference copy signals. Thus, we adjust our behavior to receive the critical information at the right time (Fooken et al., 2024; Keshava et al., 2024), and this strategic timing of behavior optimizes game performance (Hayhoe et al., 2012). The brain, in the role of a conductor, coordinates the dynamic interplay of the available movement instruments. This culminates in a pursuit solo shortly before the interception, which was most relevant for successful actions.

### Fine-tuning behavioral instruments with practice and task demands

Our results show that the orchestration of behavior is impressively consistent and efficient, with only slight adaptations to changing conditions like target speed and paddle size. This highlights our ability to optimize movements for task success, making subtle or gradual adjustments as situations change. We constantly experience a changing environment in everyday life where we need to interact with dynamic objects, allowing us to quickly tap into effective motor responses with our eyes, head, and hands. Remarkably, we demonstrated that people automatically show robust movement patterns and sustain these patterns across various conditions, with only a little bit of fine-tuning depending on experience and the current task demands.

Our findings suggest that the increased pursuit response is highly trained and automatically applied in interception games, since it was already established within the initial few actions (**see Fig. 4 and 5**). Participants concentrated on the approaching phase from the start (first 15 actions, also in Hoppe et al., 2018), and this pattern was maintained throughout the experiment and across conditions. Surprisingly, both head movement frequency and performance increased slightly over time, even though head movements were generally associated with worse performance. Potentially moving your head after longer exposure is an expression of muscular tension or a way to compensate for beginning fatigue. While pursuit frequency increased slightly for fast balls and small paddles, performance declined under these conditions, suggesting that they pose greater demands on participants. Although participants tried to compensate with closer tracking, higher pursuit rates did not fully offset performance decreases.

### Continuous tasks in computer games to study behavior

By combining mobile eye tracking with state-of-the-art computer vision, we establish a methodology that leverages games to study the coordination of eye, head, and hand movements. Our approach extends previous gamified experiments (Avraham et al., 2017; Kirsch & Kunde, 2018; Laitin & Witt, 2020; Mikula et al., 2022; Shalom et al., 2011) by focusing on the investigation of complex, sequential behaviors which points to the outstanding potential of computer games as a tool for exploring human behavior. Gamified tasks keep participants motivated, providing a natural goal that makes the task both engaging and realistic. As an experimental tool, they enable us to study dynamic, less predictable, real-world behaviors while maintaining the precision and control of experimental settings—an exciting step forward from isolated, trial-by-trial approaches.

## Conclusion

Using advanced mobile eye-tracking technology, we captured eye, head, and hand movements in a dynamic computer game with continuously changing demands. Our findings reveal how participants naturally adapt their behavior to the demands of the upcoming task— minimizing information loss by performing some movements (blinks, saccades and head movements) at non-critical times and enhancing pursuit eye movements precisely at moments critical for successful interception. This pattern of behavior was strongly linked to higher interception performance, showcasing the successful orchestration of movement patterns to meet continuously changing demands of our environment.

## Acknowledgements

We thank Jelmer de Vries for programming the first version of the game and Nils Borgerding for helping with data collection. Additionally, we thank Jolande Fooken and Philipp Kreyenmeier for helpful discussions. We acknowledge that we used AI-tools (Chat-GPT-4 and Perplexity.ai) to improve readability of some paragraphs.

## Data availability

Data and code are publicly available on OSF: [dataset] Schroeger, A., Goettker, A., Braun, D. I., & Gegenfurtner, K. R. (2025). Pong - Rapidly orchestrated eye, hand, and head movements. Open Science Framework. https://doi.org/10.17605/OSF.IO/W9VTJ

